# Beyond eplet counts: machine learning integration of evolutionary distance and electrostatic divergence improves prediction of HLA-DQ donor-specific antibody formation after kidney transplantation

**DOI:** 10.64898/2026.07.17.739230

**Authors:** Ofek Kirshenboim, Yoram Louzoun

**Author notes:** Corresponding author: Yoram Louzoun, Department of Mathematics, Bar-Ilan University, Ramat Gan, 5290002, Israel.

## Abstract

The emergence of donor-specific antibodies (DSA) against HLA-DQ reduces graft survival and limits future transplant options. Current practice predicts DSA development from HLA antigen mismatches or unique eplet counts (HLAMatchmaker), but predictive accuracy remains limited. Improving the prediction accuracy is crucial for solid organ transplants. We analysed a retrospective cohort of 240 kidney transplants (480 donor-DQ samples) in which the donor DQ allele targeted by each post-transplant DSA was identified. We compared the predictive accuracy of unique eplets (HLAMatchmaker), total eplets, amino acid mismatches, PAM genetic distance, and electrostatic mismatch score (EMS), individually and in combination. DSA emerge through the response of two donor and two patient DQ heterodimers. We evaluate the appropriate combination method of patient and donor alleles. Total eplet or amino acid mismatch counts outperformed unique eplets as used in HLAMatchmaker (AUC 0.79 vs. 0.73). Combining eplets with PAM and EMS distances in an XGBoost classifier achieved AUC 0.84, with a hazard ratio above 7. The most informative features were PAM distance and a subset of eplets distributed across the HLA-DQ alpha and beta chains. When extending prediction from a single donor DQ allele to the donor as a whole, the maximum of the two per-allele scores outperformed a probabilistic combination. The unique-eplet approach is suboptimal for predicting anti-DQ DSA in this cohort; combining total eplets with genetic and biochemical distance metrics yields substantially better prediction, essential for the reduction of DSA emergence. External validation in independent cohorts is the necessary next step.

## Introduction

Kidney transplantation is the preferred treatment for end-stage renal disease, but long-term graft survival remains substantially limited by immune-mediated injury. The human leukocyte antigen (HLA) system is the human major histocompatibility complex: a highly polymorphic set of cell-surface proteins that present peptides to T cells and that typically differ at many amino acid positions between an unrelated donor and recipient. When donor and recipient HLA sequences diverge, the recipient’s immune system can raise antibodies against the mismatched donor HLA molecules; these are termed de novo donor-specific antibodies (DSA). DSA formation is the central trigger for antibody-mediated rejection (ABMR), an immune attack on the graft mediated by these antibodies, and ABMR is the most frequent cause of late allograft (transplanted organ) loss.(Hafeez *et al*. 2023) HLA-DQ is one of several HLA loci (alongside HLA-DR and HLA-DP) routinely typed before transplantation; although HLA-DR has historically attracted most attention, converging evidence indicates that HLA-DQ mismatches are at least as immunogenic (as likely to provoke an antibody response), constituting an independent predictor of ABMR and graft failure even after adjusting for DR mismatch burden.(Das and Greenspan 2025)

Risk stratification before transplantation hinges on accurately predicting the probability that a given donor-recipient pair will generate anti-DQ DSA. From a computational standpoint, this is a binary classification problem in which the input is a pair of aligned HLA protein sequences (donor and recipient) and the output is the probability of antibody formation. Molecular mismatch load (MML) scoring was proposed as a conceptual advance over simple allele-count mismatch (a coarse feature that only records whether the donor and recipient carry the same HLA allele or not), quantifying the structural divergence between donor and recipient HLA at the level of individual amino acid configurations.(Saleem *et al*. 2022; Tambur *et al*. 2018, 2020; Tambur and Das 2023) Within MML, eplets -short, structurally clustered amino-acid motifs (each defined by 2-5 surface-exposed positions within a 3.0-3.5 Å radius on the folded HLA structure) that are thought to constitute a minimal antibody-accessible epitope - have become the dominant hand-engineered feature for immunogenicity scoring; a fixed, curated registry of several hundred such eplets exists for the HLA-DQ locus, and each donor-recipient comparison can in principle be encoded as a binary vector over this registry.(Duquesnoy and Askar 2007) The HLAMatchmaker algorithm, a rule-based, non-machine-learning tool distributed as a spreadsheet rather than as source code, enumerates *unique* eplets present in the donor sequence but absent from the recipient sequence, and clinical risk thresholds based on this single summary count have been proposed and validated across several cohorts (>17 eplets for HLA-DQ,(Wiebe *et al*. 2013) 11 eplets for combined DR and DQ,(Wiebe *et al*. 2017) and 9 eplets for DQ alone(Wiebe *et al*.2019)).

In practice, however, HLAMatchmaker predictions are imperfect: many recipients develop DSA despite low unique-eplet counts, and others remain sensitisation-free despite high counts.(Wiebe *et al*. 2013) This disconnect may reflect two conceptual limitations. First, unique-eplet scoring discards information about the aggregate antigenic burden; counting all mismatched eplets (including overlapping ones) captures more of the available signal. Second, and perhaps more fundamentally, unique-eplet counts ignore the evolutionary and biophysical dimensions of HLA divergence. T-cell help for antibody class-switching - the rate-limiting immunological bottleneck for de novo DSA production - depends on the recognition of donor-derived peptides presented by recipient MHC molecules, a process that is exquisitely sensitive to the depth of molecular divergence between donor and recipient alleles, not merely to the presence of defined surface epitopes.(Demir *et al*. 2025) Consistent with this view, Maguire et al. recently demonstrated that phylogenetic group (G1/G2), total evolutionary distance, and surface electrostatic divergence between donor and recipient DQ proteins are significant determinants of DQ immunogenicity, with cross-group mismatches carrying a substantially higher antibody risk than within-group mismatches.(Maguire *et al*. 2024)

Building on this framework, we used the well-annotated cohort of kidney transplants assembled by Maguire et al., in which the specific donor DQ allele targeted by each DSA was identified, to predict DSA formation more accurately than current methods. We systematically compared eplet-based, amino-acid (AA)-based, and biophysically informed predictors of DQ DSA formation and demonstrate that their integration in a prediction model substantially outperforms current clinical tools. SHAP (SHapley Additive exPlanations) analysis provides a quantitative decomposition of the relative predictive contributions of structural, evolutionary, and electrostatic features, offering mechanistic insight into what drives DQ immunogenicity at the molecular level. The resulting predictor is provided as an open-source tool to facilitate pre-transplant risk stratification and prospective validation.

Several computational tools have been proposed to move beyond simple eplet counting, but each has so far remained a deterministic, rule-based scoring function rather than a trained, data-driven classifier. PIRCHE-II scores immunogenicity by predicting, from the donor HLA sequence and the recipient’s HLA class II binding repertoire, which donor-derived peptides can be presented to T cells, without learning any weights from outcome data.(Tian *et al*. 2025) HLA-EMMA similarly formalises amino-acid-level mismatch counting into a standardised, userfriendly pipeline, but again applies fixed counting rules rather than a fitted model.(Kramer *et al*. 2020) Physicochemical distance metrics related to our AA mismatch and PAM250 features have previously been explored by Kosmoliaptsis et al. and Tambur, but as individual, unweighted predictors rather than combined within a supervised ensemble.(Kosmoliaptsis *et al*. 2016; Tambur 2018) The present work differs from all of these in treating DSA prediction explicitly as a supervised learning problem: rather than hand-weighting or simply summing structural, evolutionary, and electrostatic distances, we let a gradient-boosted tree ensemble learn the (non-linear, interacting) mapping from these features to DSA risk directly from outcome data, and use SHAP values post hoc to recover which features and interactions the model relies on. This learned-weighting approach is what allows PAM250 and EMS - each individually only a modest predictor - to contribute strongly once combined with eplet and zygosity information in the ensemble (see the integrated-model results below).

## Methods

### Study cohort

We analysed a retrospective cohort of 240 patients who underwent kidney transplantation between 2008 and 2017 at North-western University Hospital, described in detail by Maguire et al.(Maguire *et al*. 2024) The DQ molecule is a heterodimer assembled from one DQA1 (alpha) and one DQB1 (beta) gene product, and each person carries two DQ heterodimers (one inherited from each parent); the cohort includes full DQA1 and DQB1 sequence typing for both recipients and donors, with post-transplant antibody testing (single-antigen bead Luminex assays) performed against each donor DQ protein to determine whether the recipient developed antibodies, and if so, against which specific donor DQ heterodimer. Transplant dates and times of first positive DSA testing were recorded. The study was conducted with institutional review board approval; informed consent was obtained from all participants.(Maguire *et al*. 2024) For each donor-recipient pair, each of the two donor DQ heterodimers (DQA1-DQB1, hereafter “DQ allele”) was treated as an independent sample and compared against the recipient’s full DQ profile (both recipient alleles), yielding two comparison samples per transplant (240*×*2 = 480 candidate samples). Three samples with missing electrostatic potential data and one with a protein sequence too short for pairwise alignment were excluded computationally, leaving a final analytical dataset of 476 donor-DQ versus recipient comparison samples (Fig. 1A). Each sample was labelled as a binary outcome: positive (DSA detected against that specific donor DQ heterodimer) or negative (no DSA detected against it), which is the target variable for all classifiers described below.

**Figure 1.**
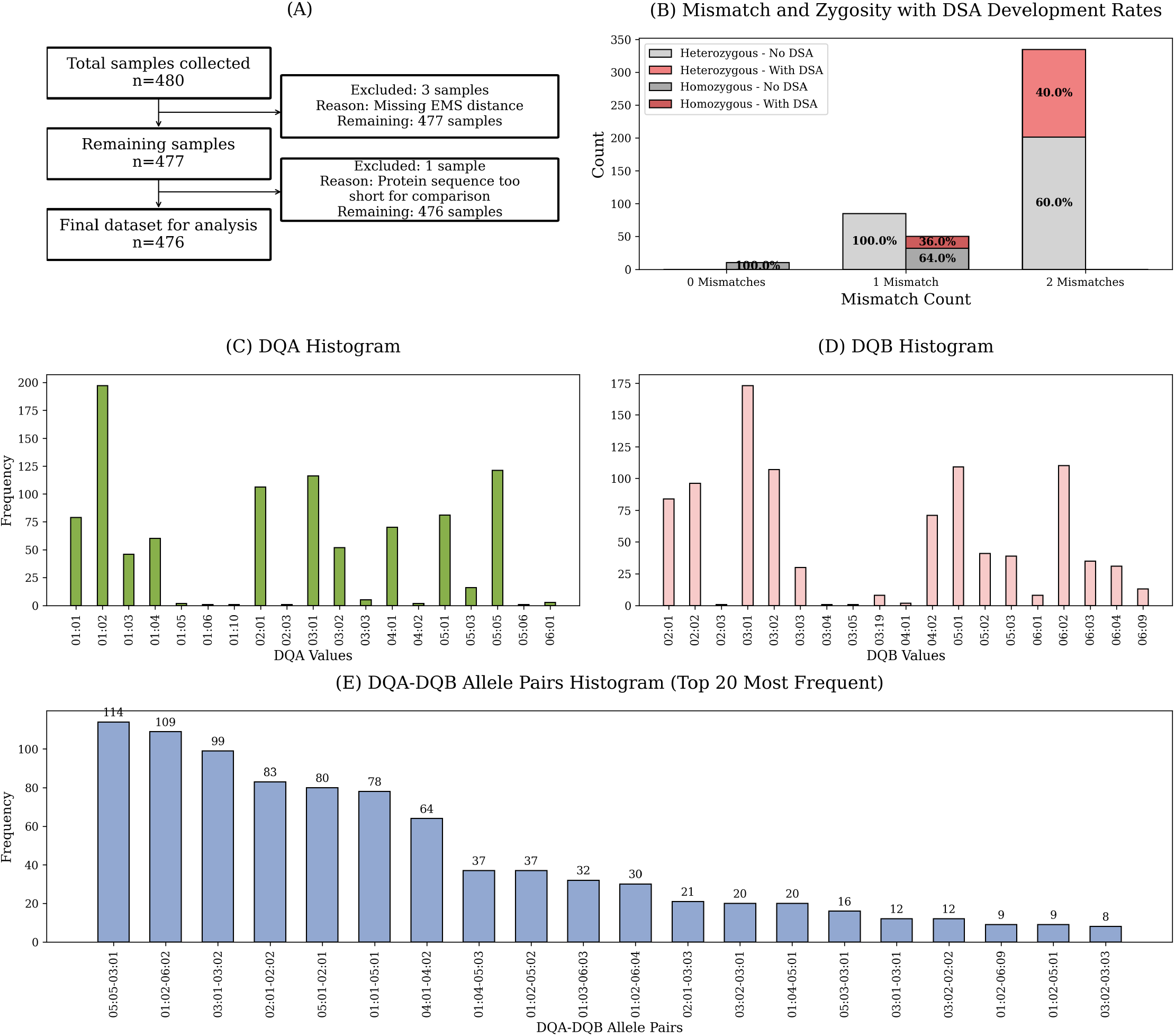
Cohort characteristics and mismatch distribution. (A) Flowchart showing sample inclusion and exclusion criteria: 480 donor-DQ versus recipient-DQ comparison samples were collected from 240 transplants; 3 samples were excluded for missing electrostatic potential data and 1 for a protein sequence too short for pairwise alignment, yielding a final analytical dataset of 476 samples. (B) Bar chart of donor-recipient DQ mismatch count (0, 1, or 2 mismatches), with stacked bars indicating the proportion of samples with (dark fill) and without (light fill) DSA development, separately for homozygous and heterozygous recipients. No DSA were detected when the donor DQ allele matched at least one recipient allele. Among fully mismatched samples, DSA rates were 36% in homozygous and 40% in heterozygous recipients. (C, D) Allele frequency histograms for DQA1 (C) and DQB1 (D) across all 476 samples; the most common alleles were DQA1*01:02 and DQA1*05:05 (C) and DQB1*03:01 and DQB1*06:02 (D). (E) Frequency histogram of the 20 most common DQA1-DQB1 heterodimer pairs; DQA1*05:05-DQB1*03:01 and DQA1*01:02-DQB1*06:02 account for the majority of samples.

### Feature engineering

#### Amino acid mismatch

All donor and recipient DQA1-DQB1 sequences in the cohort were first aligned to a common reference numbering. Polymorphic positions were identified as sites at which at least one allele in the dataset differs from the remainder; invariant positions were discarded, leaving a fixed-length vector of positions common to all comparisons. For each donor-recipient comparison, each retained position was encoded as a binary indicator (1 if the donor and recipient amino acid differ at that position, 0 otherwise), and the AA mismatch score used for prediction was the sum of these indicators (i.e. total mismatched positions) across the combined DQA1-DQB1 sequence (Fig. 2C).

**Figure 2.**
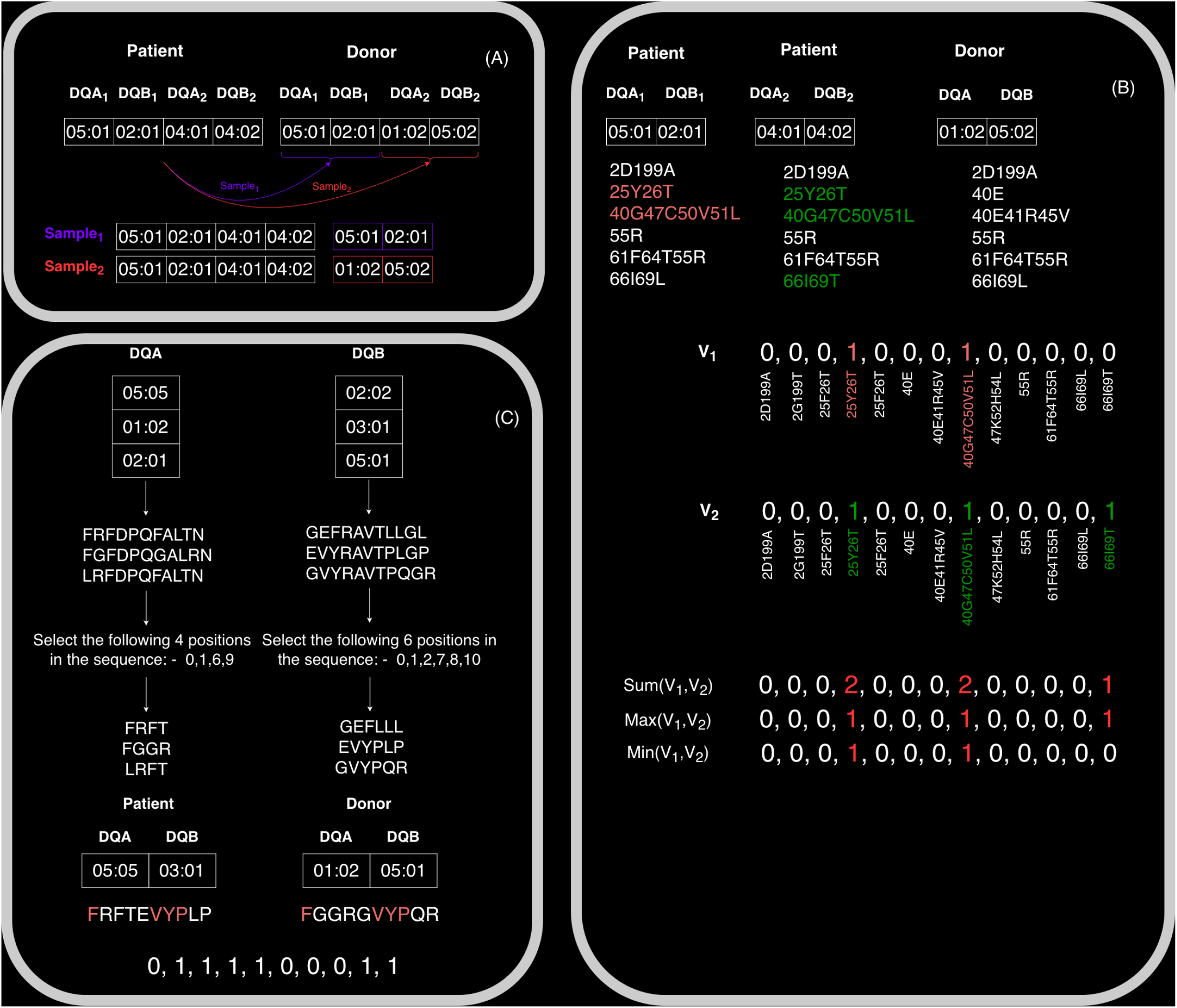
Feature engineering workflow. (A) Construction of donor-DQ versus recipient-DQ comparison samples from each transplant record. Each transplant contributes two samples, one per donor DQ heterodimer (DQA1-DQB1 pair), each compared against the recipient’s full two-allele DQ profile. (B) Eplet mismatch encoding. For a given donor allele, mismatch binary vectors *V*_1_ and *V*_2_ are computed against each of the two recipient DQ alleles (1 = eplet present in donor but absent in that recipient allele). The three aggregation rules applied across recipient alleles - sum (*V*_1_ + *V*_2_), max (element-wise maximum), and min (element-wise minimum) - are illustrated for a worked example. (C) Amino acid mismatch encoding. For the same donor-recipient pair, the mismatch score at each polymorphic position of the concatenated DQA1-DQB1 sequence is shown (1 = amino acid differs between donor and recipient allele at that position); invariant positions are excluded.

#### Eplet mismatch

Donor and recipient alleles were mapped to their amino acid sequences using the IPD-IMGT/HLA database.(Barker *et al*. 2023) A positional offset of 22 residues (DQA1) and 31 residues (DQB1) was applied to align eplet registry positions to IMGT numbering. Each donor-recipient pairing was then encoded, over the full curated eplet registry, as a fixed-length one-hot binary vector in which each entry corresponds to one registry eplet (1 = eplet present in the donor sequence but absent from the recipient sequence at the same registry position, 0 otherwise) (Fig. 2B). Two summary scores were derived from this vector: the “all-eplet” score, which sums over every entry regardless of whether the eplet has documented antibody reactivity, and the “confirmed-eplet” score, which sums only over the subset of eplets for which antibody reactivity has been experimentally confirmed in the HLAMatchmaker registry. As an external, clinically used comparator, unique eplets were separately computed with the (non-machinelearning, spreadsheet-based) HLAMatchmaker tool to allow direct comparison with the current clinical standard.(Duquesnoy and Askar 2007; Duquesnoy and Marrari 2002; Duquesnoy 2008; Duquesnoy and Marrari 2009)

#### Electrostatic mismatch score (EMS)

The three-dimensional electrostatic potential surrounding each HLA structure, derived from homology models of the folded DQ heterodimer, was computed by solving the linearised Poisson-Boltzmann equation on a grid representation of the protein surface; pairwise comparison between donor and recipient surfaces was performed using the PIPSA (Protein Interaction Property Similarity Analysis) software, which returns a single scalar distance per donor-recipient pair.(Mallon *et al*. 2018) This scalar EMS value quantifies surface charge divergence between donor and recipient alleles and has been validated as a biophysical measure of immunogenic distance by Mallon et al.(Mallon *et al*. 2018)

#### Phylogenetic distance (PAM250)

As a complementary, purely sequence-based distance metric, molecular phylogenetic analysis was performed in MEGA v11.0.13(Tamura *et al*. 2021) using maximum-likelihood estimation and the PAM250 (point accepted mutation) amino acid substitution matrix, which scores pairwise sequence divergence in terms of the log-odds of one amino acid being substituted by another over evolutionary time, summed over the aligned DQA1-DQB1 sequence to give one PAM250 distance per donor-recipient allele pair.(Dayhoff *et al*. 1978) The resulting phylogenetic tree over all observed DQ alleles resolved into two principal clades, corresponding to the presence or absence of the DQA1*01 allele family; these two clades are denoted G1 (evolutionary group 1) and G2 (evolutionary group 2), following the nomenclature of Maguire et al., and were used both as a categorical feature (G1/G2 membership of donor and recipient) and, in continuous form, as the PAM250 distance itself.(Maguire *et al*. 2024)

#### Mismatch aggregation across recipient alleles

Given a single donor DQ allele and two recipient DQ alleles, mismatch vectors *v*_1_ and *v*_2_ (one per recipient allele) were combined using three aggregation rules:

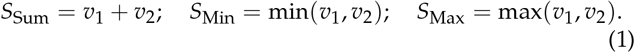

To translate allele-level scores into a patient-level risk estimate, we evaluated three combination rules given allele-level scores *S*_1_ and *S*_2_:

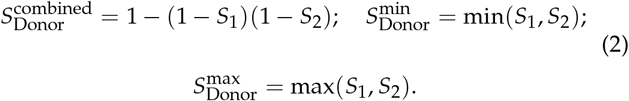

### Machine learning

Gradient-boosted decision tree classifiers, XGBoost(Chen and Guestrin 2016) and LightGBM(Ke *et al*. 2017), were implemented in Python and trained on tabular feature matrices constructed by concatenating: AA mismatch indicators, eplet mismatch vectors (all-eplet or confirmed-eplet subsets), the scalar EMS value, the scalar PAM250 distance, a categorical G1/G2 donor-recipient configuration feature, and a binary zygosity indicator (whether the recipient is homozygous or heterozygous at the mismatched DQ locus). Gradient-boosted trees were chosen over deep-learning architectures because the feature matrix is lowdimensional (tens to low hundreds of largely binary or scalar features) and highly sparse for the eplet vectors, a regime in which tree ensembles typically outperform neural networks and additionally provide native SHAP-based interpretability. Eplet and AA feature sets were not combined into a single model because their information content overlaps substantially (most AA mismatches at eplet-registry positions are, by construction, also eplet mismatches), which would otherwise inflate the apparent importance of this shared signal. The dataset was split into five stratified 80%/20% train/test partitions (5-fold crossvalidation), with donors, rather than individual samples, assigned consistently to either the training or the test partition in every fold, so that both DQ alleles of a given donor were always kept together; this donor-level splitting prevents information leakage that would arise if the two alleles of the same donor were separated across the training and test sets. Standard hyperparameters were used without tuning (XGBoost: 100 estimators, learning rate 0.3, default tree depth; LightGBM: 100 estimators, learning rate 0.1, default tree depth); no hyperparameter search was performed, since the goal was to compare feature sets rather than to optimise absolute performance, and given the limited size of the cohort. Feature importance was characterised using SHAP (SHapley Additive exPlanations) values, which decompose each individual prediction into additive per-feature contributions.(Lundberg and Lee 2017) Models were built iteratively over increasing feature subsets (features added in decreasing order of training-set mean absolute SHAP importance, in increments of two) to identify the feature count at which test-set performance plateaued, i.e. a data-driven feature-selection curve rather than a fixed feature count chosen a priori. We also evaluated dedicated time-to-event models on the same feature sets, since DSA formation is technically a right-censored event (patients without DSA at last follow-up may still develop it later): a Random Survival Forest (RSF)(Ishwaran *et al*. 2008), which extends random forests to censored outcomes via a log-rank splitting criterion, and an XGBoost classifier with a Tobit-style censoring-aware loss that explicitly models the censoring time rather than treating DSA-free follow-up as a negative label.(Barnwal *et al*. 2022; Shtossel *et al*. 2025) Both were trained under the same five-fold, donor-level cross-validation scheme described above, with concordance index (CI) as the evaluation metric.

### Statistical analysis and evaluation

Model discrimination was assessed by the area under the receiver operating characteristic curve (AUC-ROC), computed using scikit-learn.(Pedregosa *et al*. 2011) To relate each continuous predictor to clinical time-to-event outcomes, patients were dichotomised at a single threshold (the point on the ROC curve closest to the top-left corner, i.e. the operating point jointly maximising sensitivity and specificity) into predicted high-risk and low-risk groups, and Kaplan-Meier curves (non-parametric estimates of the fraction of patients remaining DSA-free over time) were compared between the two groups using the log-rank test; the hazard ratio (HR) is the corresponding relative rate of DSA formation in the high-risk versus the low-risk group; a HR of 1 indicates no difference in risk, and larger values indicate stronger clinical separation. Predictive accuracy for time-to-DSA was additionally quantified without dichotomisation by the concordance index (CI, the probability that, for a random pair of patients, the model ranks the one who developed DSA sooner as higher risk), which is the standard discrimination metric for survival models and is analogous to the AUC but accounts for censoring.(Harrell *et al*. 1982) The association between G1/G2 donor-recipient configurations and DSA development was assessed with a *χ*^2^ test of independence on the corresponding contingency table. All analyses were performed in Python (SciPy, Matplotlib). Standard errors and 95% confidence intervals on AUC values were derived analytically using the Hanley & McNeil (1982) approximation for the variance of the Wilcoxon (Mann-Whitney U) statistic, avoiding the need for computationally expensive bootstrapping.(Hanley and McNeil 1982)

## Results

### Cohort characteristics and mismatch distribution

To ensure a direct association between mismatch and DSA, we focused on a single-centre design in which the association of DSA with each donor DQ heterodimer is known. The analytical dataset comprised 476 donor-DQ versus recipient comparison samples derived from 240 transplants (4 samples were removed - Fig. 1A). No DSA were detected in any transplant where the donor DQ allele exactly matched at least one of the recipient’s two DQ alleles, confirming that a full sequence mismatch (i.e. the donor DQ heterodimer differing from both recipient DQ heterodimers) is a necessary condition for sensitisation (antibody formation) in this cohort. Among fully mismatched samples, DSA were detected in 36% of samples from recipients who are homozygous at the DQ locus (i.e. the recipient’s two DQ alleles are identical to each other, so the donor allele is mismatched against a single distinct recipient sequence) and in 40% of samples from heterozygous recipients with two mismatches (i.e. the donor allele differs from both of the recipient’s two distinct DQ alleles) (Fig. 1B). The most common DQB1 alleles were DQB1*03:01 and DQB1*06:02; the most common DQA1 alleles were DQA1*01:02 and DQA1*05:05 (Fig. 1C,D). The most frequent DQ heterodimer pairs, DQA1*05:05-DQB1*03:01 and DQA1*01:02-DQB1*06:02, accounted for the majority of samples (Fig. 1E).

### Eplet and amino acid mismatch as individual predictors of anti-DQ DSA

We analysed each donor DQ heterodimer individually and computed the association between AA mismatch and eplet mismatches and DSA development against each donor DQ het-erodimer (see description of the analysis in Fig. 2).

Both eplet and AA mismatch showed a clear monotonic relationship between the sum of mismatches of the donor with both recipient alleles and DSA probability (Fig. 3A,B). Because each donor DQ heterodimer can have a different mismatch count with each of the two recipient DQ heterodimers, we tested three aggregation strategies (Min, Max, and Sum; Eq. 1) and computed the AUC of aggregated counts against DSA emergence. *S*_Sum_yielded the highest AUC for eplets (*p* = 0.0004 vs *S*_Min_), whereas *S*_Min_ was optimal for AA mismatch (*p* = 0.0005 vs *S*_Max_). Critically, all-eplet counts (AUC 0.79) substantially outperformed HLAMatchmaker unique eplets (AUC 0.73), and direct AA mismatch performed comparably to all-eplet counts (AUC 0.78) (see Fig. 3C-H for all ROC and Kaplan-Meier curves). Even the coarsest possible predictor - a binary matched/mismatched classification - achieved AUC 0.645 (Supp. Mat. Fig. S1), which means HLAMatchmaker unique-eplet scoring captures approximately half the available discrimination between a match/mismatch baseline and full eplet or AA-level scoring.

**Figure 3.**
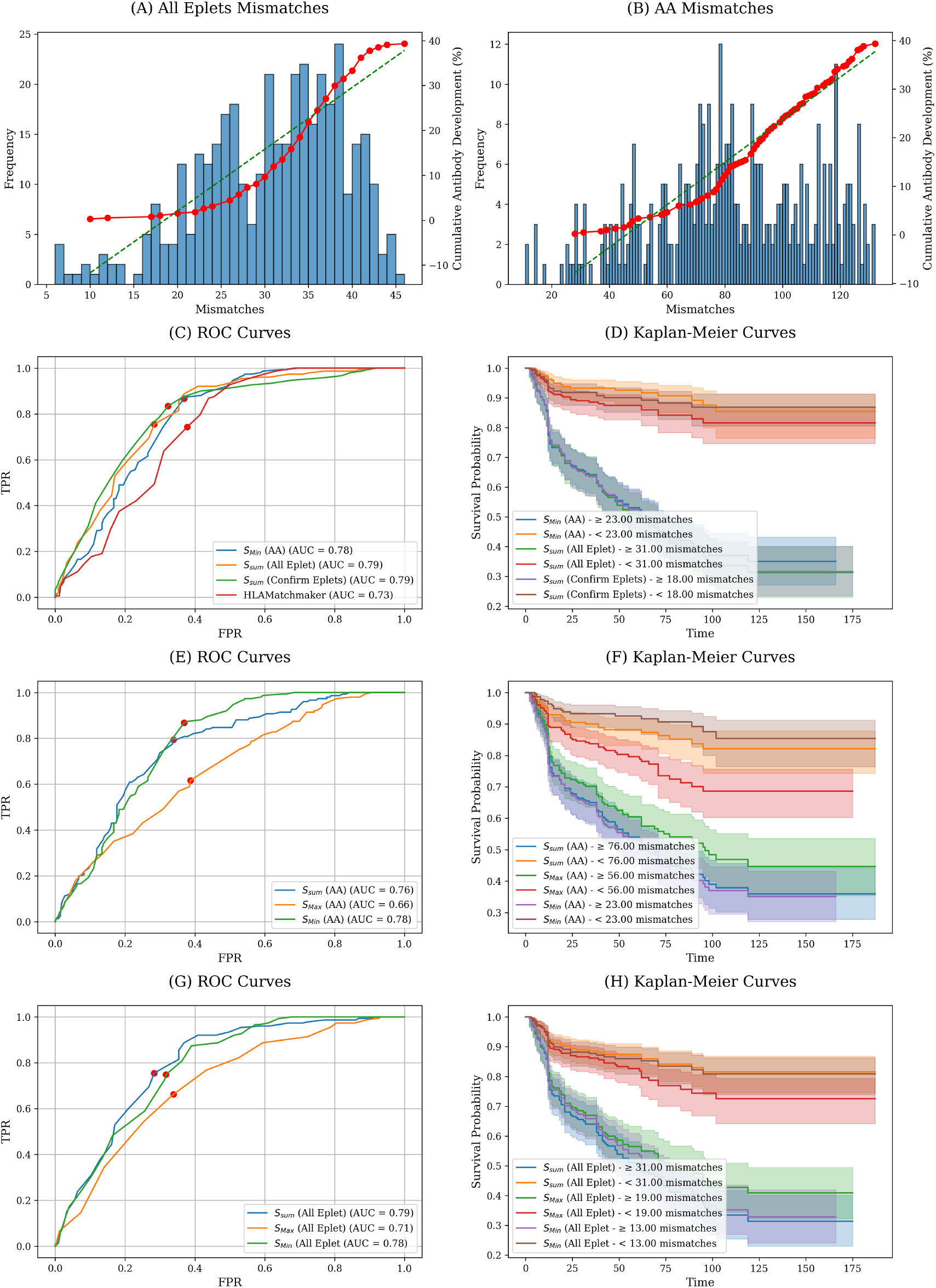
Performance of eplet and amino acid mismatch as individual predictors of anti-DQ DSA formation. (A, B) Relationship between mismatch count and DSA development for all-eplet mismatches (A) and amino acid (AA) mismatches (B). Bar height shows the frequency of samples at each mismatch count; the red curve shows the cumulative DSA development rate; the green dashed line shows a linear fit. Both predictors show a monotonic relationship with DSA probability. (C) ROC curves comparing four individual predictors at the allele level, using the aggregation strategy that maximised AUC for each: AA mismatch with *S*_Min_ (AUC 0.78), all-eplet count with *S*_Sum_ (AUC 0.79), confirmed-eplet count with *S*_Sum_ (AUC 0.79), and HLAMatchmaker unique eplets (AUC 0.73). Red dots indicate the point of optimal accuracy (closest to the top-left corner). (D) Kaplan-Meier survival curves (time to DSA formation) for the three main predictor types at their optimal accuracy thresholds; threshold values are shown in the legend. Shaded regions indicate 95% confidence intervals. (E, F) Sensitivity analysis of aggregation strategy (*S*_Sum_, *S*_Max_, *S*_Min_) for AA mismatch: ROC curves (E) and Kaplan-Meier curves at the optimal threshold for each aggregation (F). (G, H) Sensitivity analysis of aggregation strategy for all-eplet mismatch: ROC curves (G) and Kaplan-Meier curves (H).

Kaplan-Meier analysis at optimal accuracy thresholds demon-strated meaningful clinical stratification: HR 4.66 for all eplets (threshold: 31 mismatches), HR 6.26 for confirmed eplets (threshold: 18 mismatches), and HR 7.14 for AA mismatch (threshold: 23 positions). Concordance indices ranged from 0.70 to 0.72 across all three feature sets; these values, obtained using AA or total eplet mismatches, exceed those achieved with the unique-mismatch approach of HLAMatchmaker.

In summary, when comparing a set of recipient DQ heterodimers against a single donor DQ heterodimer, taking the sum of all eplet differences across both recipient alleles yields the best discrimination. No difference was observed between models using only confirmed eplets and those using all eplets.(Senev *et al*. 2020)

### Evolutionary and electrostatic features independently predict DSA formation

Beyond eplet and AA counts, we evaluated three evolutionary and biophysical features: (i) G1/G2 phylogenetic category; (ii) PAM250 genetic distance; and (iii) EMS. G1-G2 cross-group transplants (G1*→* G2/G2 or G2*→* G1/G1) carried a 50% DSA risk, compared with 11-18% for same-group homozygous transplants. The lowest-risk configuration was an G1 donor to an G1/G2 heterozygous recipient (7%); an G2 donor to a heterozygous recipient carried 40% risk (Fig. 4A). EMS significantly stratified DSA risk in Kaplan-Meier analysis, with an AUC of 0.62, but PAM250 distance alone did not reach significance (AUC 0.55) (Fig. 4B-D for AUC, hazard ratio, and CI; Fig. 4E-H for ROC and Kaplan-Meier curves across all aggregation strategies).

**Figure 4.**
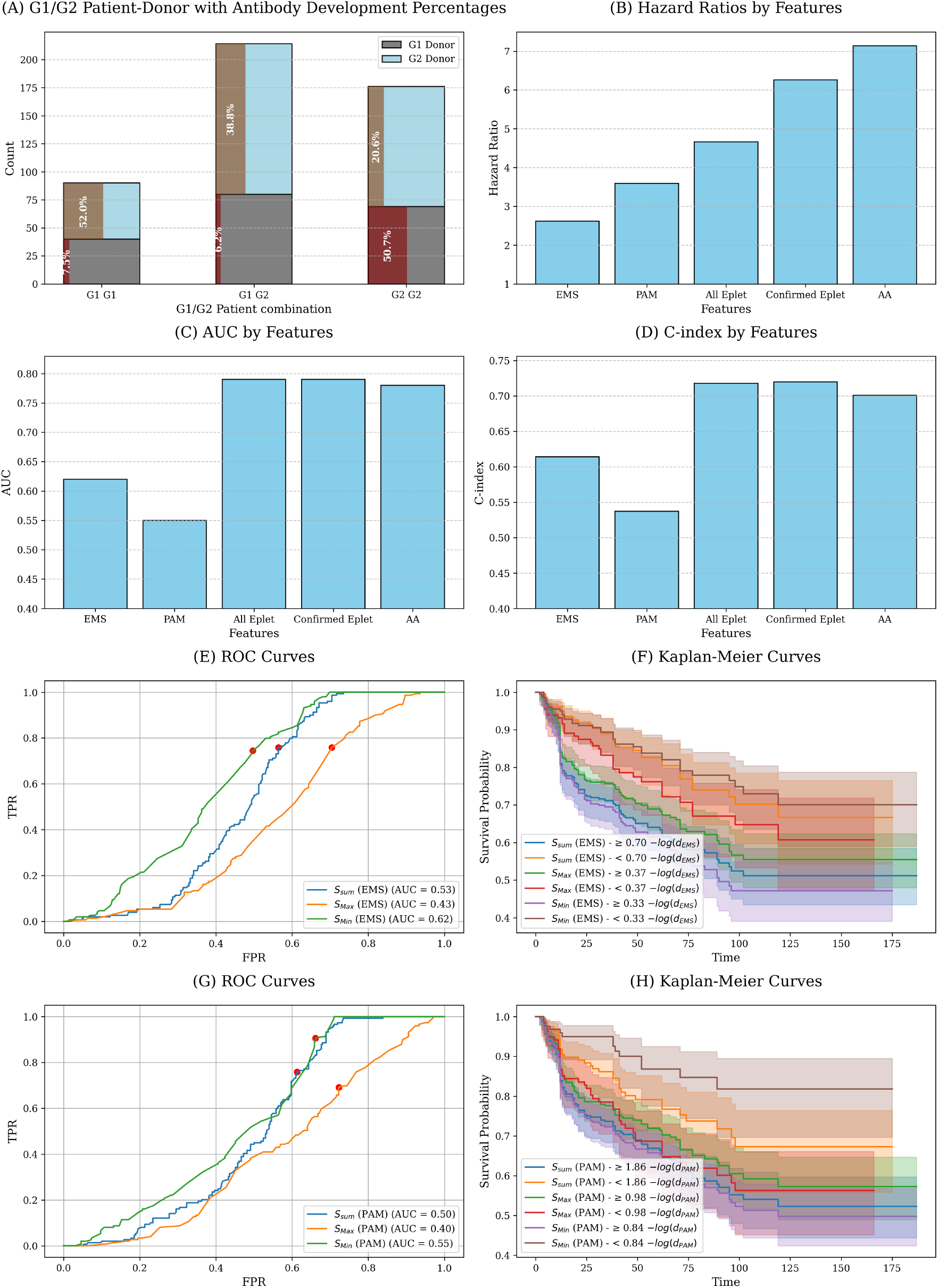
Evolutionary and electrostatic features as predictors of DSA formation. (A) DSA development rates stratified by G1/G2 donor-recipient configuration. Bars show the number of samples in each configuration (G1 *→*G1, G1*→* G2, G2*→* G1, G2*→* G2, where the arrow indicates donor *→*recipient genotype); the percentage within each bar indicates the DSA rate. Cross-group transplants (G1*→* G2 or G2*→* G1) carried approximately 50% DSA risk, compared with 11-18% for same-group configurations. (B) Hazard ratios (at the optimal accuracy threshold) for each feature set - EMS, PAM250, all-eplet, confirmed-eplet, and AA mismatch - using the aggregation strategy that maximised AUC for each. (C, D) Performance of electrostatic mismatch score (EMS) across the three aggregation strategies: ROC curves (C) and Kaplan-Meier survival curves at the optimal threshold for each aggregation (D). Shaded regions indicate 95% confidence intervals. (E, F) Performance of PAM250 phylogenetic distance across the three aggregation strategies: ROC curves (E) and Kaplan-Meier curves (F).

### Integrated machine learning model improves stratification

Combining all features of each donor DQ heterodimer in an XGBoost classifier and ranking by SHAP importance (Fig. 5C), performance plateaued at 20-30 features, beyond which additional features did not improve discrimination (Fig. 5D). The most influential features were PAM250 distances for each DQA1DQB1 pair (across both recipient alleles), followed by EMS values, AA mismatch scores, and a limited panel of specific eplets; the G1/G2 binary category was not among the top 20 features, suggesting that the categorical grouping captures only a fraction of the biologically relevant information encoded in the continuous evolutionary distance.

**Figure 5.**
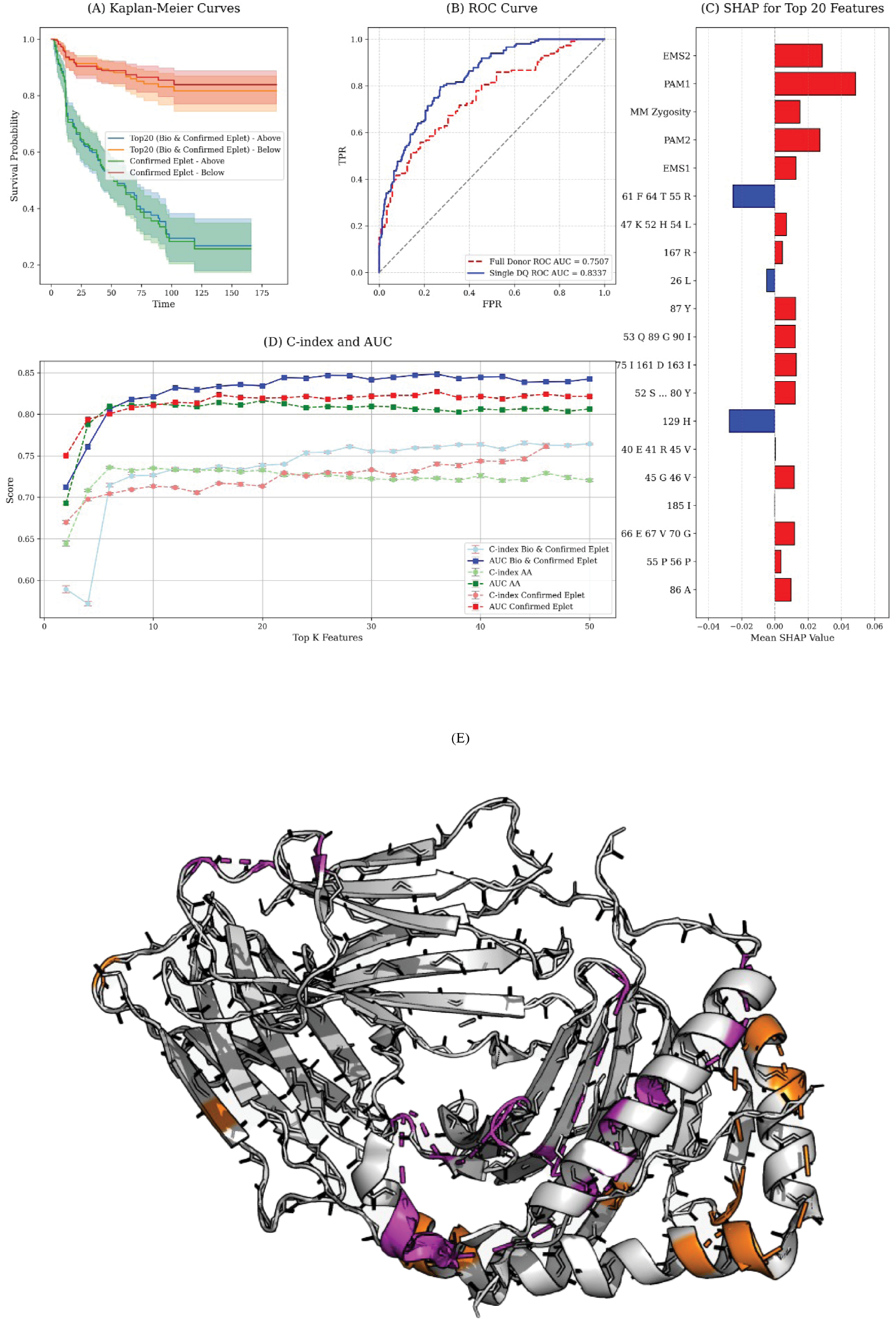
Integrated machine learning model. (A) Kaplan-Meier survival curves (time to DSA formation) for the integrated XG-Boost model (top 20 biophysical + confirmed-eplet features) versus confirmed-eplet count alone, at the respective optimal accuracy thresholds. (B) ROC curves comparing allele-level prediction (single-donor DQ heterodimer; AUC 0.8337) and full-donor prediction (AUC 0.7597). (C) Mean absolute SHAP values for the top 20 features in the integrated XGBoost model, ordered by importance. EMS and PAM250 distances (computed against each of the two recipient alleles) are the dominant contributors. (D) AUC (filled symbols) and concordance index (open symbols) as a function of the number of features included (top-k analysis), for three featureset strategies: biophysical + confirmed-eplet features (circles), AA mismatch features (squares), and confirmed-eplet features alone (triangles). (E) Structural localisation of the informative eplets from the top-20 feature set. Residues are highlighted in magenta (alpha chain), and orange (beta chain). Informative positions span both chains and both alpha-helical and loop regions, consistent with a distributed immunogenic surface.

The final integrated model achieved AUC 0.84 and CI 0.82, versus AUC 0.79 for the best single-feature predictor and AUC 0.73 for HLAMatchmaker unique eplets. Table 1 summarises the performance of all feature sets. Kaplan-Meier analysis confirmed improved risk stratification compared with single-feature models (Fig. 5A). Survival forest and censoring-aware gradient-boosted (Tobit-loss) time-to-event models were also evaluated but did not exceed the XGBoost performance reported here (Supp. Mat. Fig. S2).

**Table 1.** Prediction performance summary across feature sets and the integrated model. AUC: area under the ROC curve; CI: concordance index; HR: Kaplan-Meier hazard ratio at the optimal accuracy threshold; EMS: electrostatic mismatch score; PAM: PAM250 phylogenetic distance; HLAMm: HLAMatch-maker unique eplets. The integrated XGBoost model combines all feature types. *For the binary match/mismatch predictor the hazard ratio is undefined (infinite) because no DSA occurred in the matched group.

| Feature set | AUC | CI | HR |
| --- | --- | --- | --- |
| HLAMatchmaker unique eplets (HLAMm) | 0.73 | 0.66 | 4.24 |
| Binary match/mismatch | 0.64 | 0.61 | ∞* |
| AA mismatch ($S_{\text{Min}}$) | 0.78 | 0.70 | 7.14 |
| All eplets ($S_{\text{Sum}}$) | 0.79 | 0.72 | 4.66 |
| Confirmed eplets ($S_{\text{Sum}}$) | 0.79 | 0.71 | 6.26 |
| EMS ($S_{\text{Sum}}$) | 0.62 | 0.61 | 2.62 |
| PAM250 ($S_{\text{Sum}}$) | 0.55 | 0.53 | 3.59 |
| **Integrated XGBoost (top 20 features)** | **0.84** | **0.82** | 7.20 |
| Full-donor prediction ($S_{\text{max}}$) | 0.76 | 0.68 | 6.94 |

Informative eplets were distributed across both the alpha (magenta) and beta (orange) chains and spanned both alphahelical and loop regions of the HLA-DQ heterodimer (Fig. 5E), indicating that the model captures a broad immunogenic surface rather than a single restricted binding pocket.

### Donor-level prediction

The clinically relevant outcome is whether a recipient will develop DSA against *any* allele of the donor DQ. Of the three combination rules (Eq. 2), the maximum allele-level score 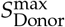was the strongest patient-level predictor (Fig. 5B). This is consistent with a censoring explanation, i.e. that the two per-allele outcomes for a given donor are not statistically independent observations but are truncated by a shared clinical event: the observed probability that a patient who developed one DSA also developed a second DSA against the same donor’s other allele (74.58%) was substantially lower than the value expected if the two allele-level events were statistically independent (82.75%, i.e. the product of the two marginal probabilities scaled appropriately), suggesting that once a DSA forms against one allele, the patient is removed from further observation - through graft failure or re-transplantation - before a response to the second allele has the opportunity to develop and be recorded (see also score correlation in Supp. Mat. Figure S3-A). There was no difference between the machine learning model trained on confirmed eplets and that trained on all eplets (Fig. 5A). Beyond discrimination (AUC, CI), we assessed whether the predicted probabilities themselves are clinically usable risk estimates by comparing predicted risk against observed DSA frequency within predicted-risk bins; both the single-DQ allele-level model and the full-donor model are well calibrated, with observed event rates tracking predicted risk closely across the probability range (Supp. Mat. Fig. S3-B,C). This calibration check is a necessary complement to AUC/CI, since a model can rank patients correctly (high AUC) while still producing systematically over- or under-confident absolute risk estimates that would be misleading if used directly for clinical decision-making.

## Discussion

The central finding of this study is that integrating molecular, evolutionary, and biophysical features of HLA-DQ mismatch in a gradient-boosted classifier achieves substantially higher predictive accuracy (AUC 0.84, CI 0.82) for de novo anti-DQ DSA formation than any single feature type, and markedly outperforms the clinical standard HLAMatchmaker (AUC 0.73). This improvement is not merely statistical: an AUC increase from 0.73 to 0.84 represents a meaningful gain in clinical discriminability that could translate into better pre-transplant risk stratification and more informed donor selection.

A key finding with immediate practical implications is that counting *all* mismatched eplets (AUC 0.79), rather than only the *unique* eplets reported by HLAMatchmaker, already provides meaningful improvement over the clinical standard, and this change requires no additional computation beyond what HLA-Matchmaker already provides. Furthermore, simple AA mismatch counts at polymorphic positions perform at the same level as all-eplet counts - a result consistent with Kosmoliaptsis et al. (2016)(Kosmoliaptsis *et al*. 2016) and Tambur et al. (2018)(Tambur 2018), who questioned the assumption of uniform immunogenicity across all eplets, and with Callemeyn et al.(Callemeyn *et al*. 2022) who observed inconsistent DSA development despite high eplet mismatch loads. Together, these results suggest that the structural framing of eplets does not capture immunogenicity information beyond what is already present in the total sequence divergence between donor and recipient.

The most consequential finding from SHAP analysis is that PAM250 evolutionary distance and EMS electrostatic divergence together constitute the dominant predictors in the integrated model. This points to a mechanistically coherent explanation: antibody production by B cells against a donor HLA protein requires help from CD4 T cells, which is triggered when donor-derived peptide fragments of the mismatched HLA protein are themselves displayed (presented) on the recipient’s own MHC class II molecules; the efficiency of this presentation step is sensitive to the global sequence divergence between donor and recipient alleles, which is precisely what the PAM250 and EMS distance metrics quantify computationally.(Demir *et al*. 2025)

Although PAM250 and EMS show modest discrimination individually, their non-linear interactions with eplet profiles and zygosity in the ensemble model substantially amplify their contribution, as reflected by their dominant SHAP importance. Alleles that are deeply divergent - as measured by PAM250 or by surface electrostatic dissimilarity - generate more foreign-looking peptide repertoires and stronger T-cell help, independently of whether those differences happen to map onto catalogued eplet positions.

The G1/G2 binary classification, which captures the most salient evolutionary split in DQ diversity, was clinically meaningful at the group level but contributed less than the continuous PAM250 and EMS measures once all features were modelled jointly. This implies that the categorical grouping is a coarse proxy for a continuously varying underlying signal, and that the full predictive value of evolutionary divergence is better captured by continuous metrics. This finding also illustrates a broader point about the choice of computational approach: fixed-rule scores such as PIRCHE-II and HLA-EMMA(Tian *et al*. 2025; Kramer *et al*. 2020) address partially overlapping aspects of HLA divergence to our AA and eplet features, but, being deterministic functions of the sequence data rather than fitted models, they cannot re-weight or discover interactions between features the way a trained ensemble can; the G1/G2 result is a concrete example of a feature (categorical evolutionary group) that looks informative in isolation but is subsumed by a continuous feature (PAM250 distance) once the two are allowed to compete for importance within a single model. Direct integration of PIRCHE-II- or HLA-EMMA-derived scores as additional input features to our XGBoost pipeline, rather than as separate scoring systems, is a natural extension and may be complementary to the present framework; their integration with biophysical distance metrics warrants investigation, ideally on a cohort large enough to support the additional features without overfitting.

At the patient level, the observation that the maximum allele-level score outperforms the probabilistic combination rule has a straightforward clinical interpretation: the relevant clinical event is DSA development against the most immunogenic donor allele, and once that event occurs (or causes graft failure), the second allele is no longer under observation. This censoring pattern is supported by the quantitative discordance between the observed and independence-expected rates of dual-allele responses. For clinical risk stratification, the practical implication is that the pre-transplant risk score for a given donor should reflect the more divergent DQ allele, not the average or sum of both.

The informative eplets identified by the model were distributed across both the alpha and beta chains of the HLA-DQ heterodimer and spanned both alpha-helical and loop regions. The absence of a single dominant immunogenic hotspot is consistent with the whole-molecule biophysical metrics (EMS, PAM250) outperforming region-specific eplet catalogues, and suggests that the immune response to HLA-DQ mismatch reflects global surface divergence rather than a restricted set of contact residues.

This study has several limitations. The cohort derives from a single centre and, being a reanalysis of data from Maguire et al., the model has not been tested in an independent external dataset; calibration was assessed only within this single cohort (Supp. Mat. Fig. S3-B,C), and calibration is not guaranteed to transfer to populations with a different DSA base rate or HLA allele frequency distribution, so recalibration (e.g. Platt scaling or isotonic regression on a local validation set) would likely be required before deployment elsewhere. External validation in multicentre prospective cohorts is an essential next step before clinical deployment, and we provide the full open-source code (https://github.com/Ofekirsh/dq-antibody-predictor) to facilitate this. The analysis is restricted to HLA-DQ; extension to HLA-DR and HLA-DP would broaden clinical applicability. The model is observational, and the relative contributions of immunosuppression regimen, prior sensitisation history, and T-cell alloreactivity to DSA formation cannot be fully disentangled from HLA mismatch effects.(Thaunat *et al*. 2016; Aubert *et al*. 2017; Kosmoliaptsis *et al*. 2014)

## Key Points

- Recasting HLA-DQ immunogenicity prediction as a supervised learning problem - rather than a fixed, rule-based scoring function as in HLAMatchmaker, PIRCHE-II, or HLAEMMA - lets a gradient-boosted tree ensemble learn non-linear interactions between structural, evolutionary, and electrostatic features directly from outcome data.
- Counting all mismatched eplets, rather than only the unique eplets used by HLAMatchmaker, already improves prediction of anti-DQ DSA (AUC 0.79 vs. 0.73) at no extra computational cost.
- PAM250 evolutionary distance and electrostatic mismatch score (EMS) are the dominant contributors in an integrated gradient-boosted model, outperforming eplet- and aminoacid-based scores alone.
- An XGBoost classifier combining eplet, amino acid, evolutionary, and electrostatic features achieves AUC 0.84 and concordance index 0.82, with a hazard ratio above 7, and remains well calibrated within the study cohort.
- At the donor level, the maximum of the two per-allele risk scores is the strongest predictor of DSA development against either donor DQ allele.

## Abbreviations

ABMR: antibody-mediated rejection
AUC: area under the receiver operating characteristic curve
CI: concordance index
DSA: donor-specific antibody
EG: evolutionary group
EMS: electrostatic mismatch score
HLA: human leukocyte antigen
MML: molecular mismatch load
PAM: point accepted mutation
SHAP: Shapley additive explanations.

## Data Availability Statement

The data analysed in this study were obtained from the cohort described in Maguire et al. (HLA, 2024; PubMed ID 38575370). The dataset is not publicly available; access requests should be directed to the authors of the original study. All code implementing the machine learning predictor described here is publicly available at https://github.com/Ofekirsh/dq-antibody-predictor.

## Ethics Statement

The cohort analysed in this study was previously collected under the approval of the Northwestern University institutional review board, with informed consent obtained from all participants, as described in Maguire et al. (HLA, 2024). The present analysis used de-identified data from that cohort and required no additional ethics approval.

## Acknowledgments

The authors thank the team responsible for the original cohort assembly and data curation at Northwestern University.

## Author Biographies

**Yoram Louzoun**. Yoram Louzoun is a Professor of Mathematics at the Bar Ilan University in Israel. He has graduated from the Hebrew University in 2000 and continued to a post-doc in Princeton. In 2002, he started a position at Bar Ilan in the Mathe-matics department. He studies stochastic processes and machine learning, with a special focus on its application to immunology, transplants and microbiome. He has developed a set of algorithms to optimize Stem cells transplants, and machine learning algorithms to predict the host condition from his microbiome or from his T cell repertoire, as well as graph based algorithms.

**Ofek Kirshenboim**. Ofek Kirshenboim performed the data analysis and implemented the machine learning pipeline described in this manuscript, and is affiliated with the Department of Mathematics at Bar-Ilan University.

## Supplementary Material Figures

**Figure S1.**
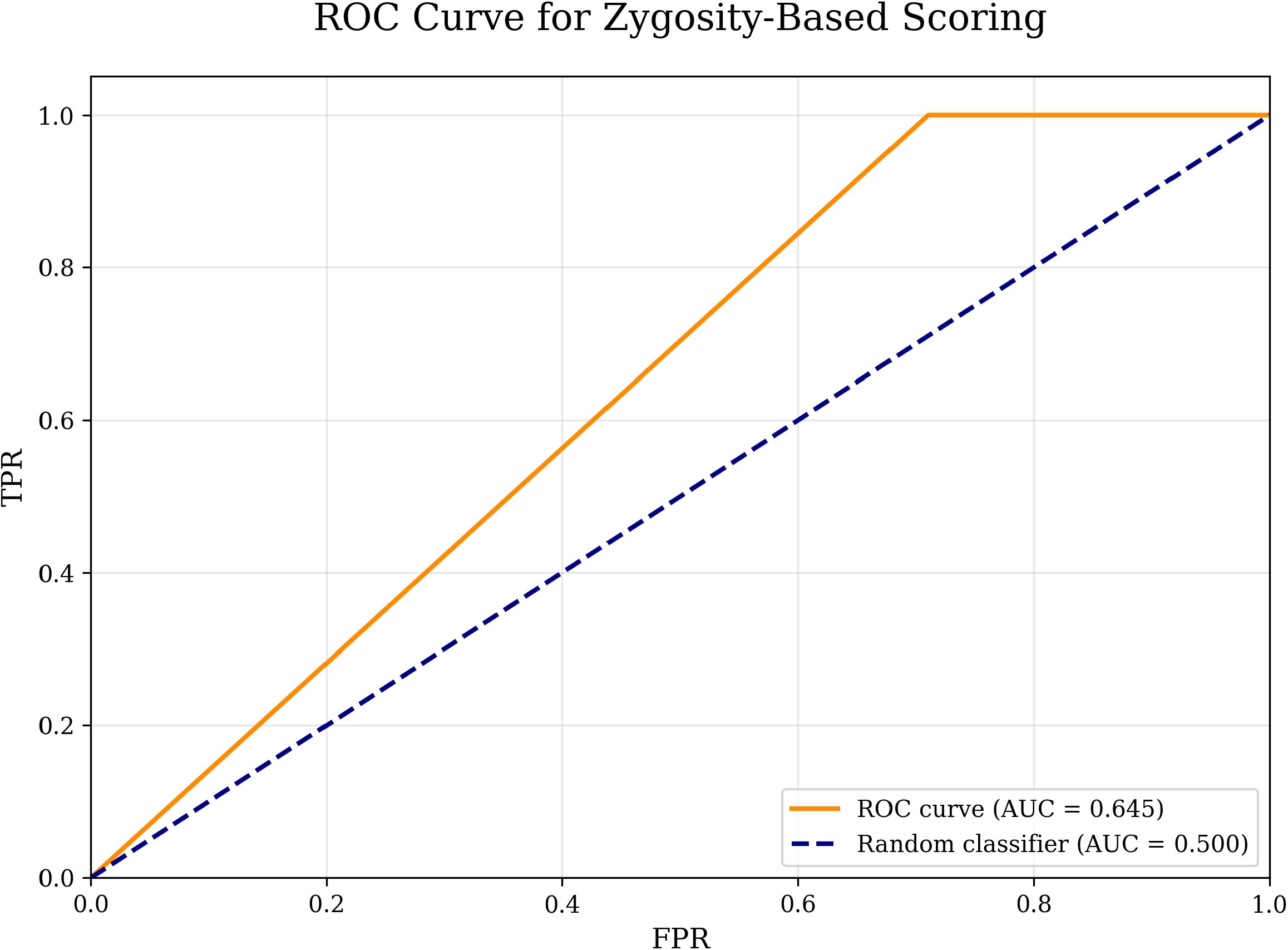
ROC curve of DSA prediction using only the match/mismatch information.

**Figure S2.**
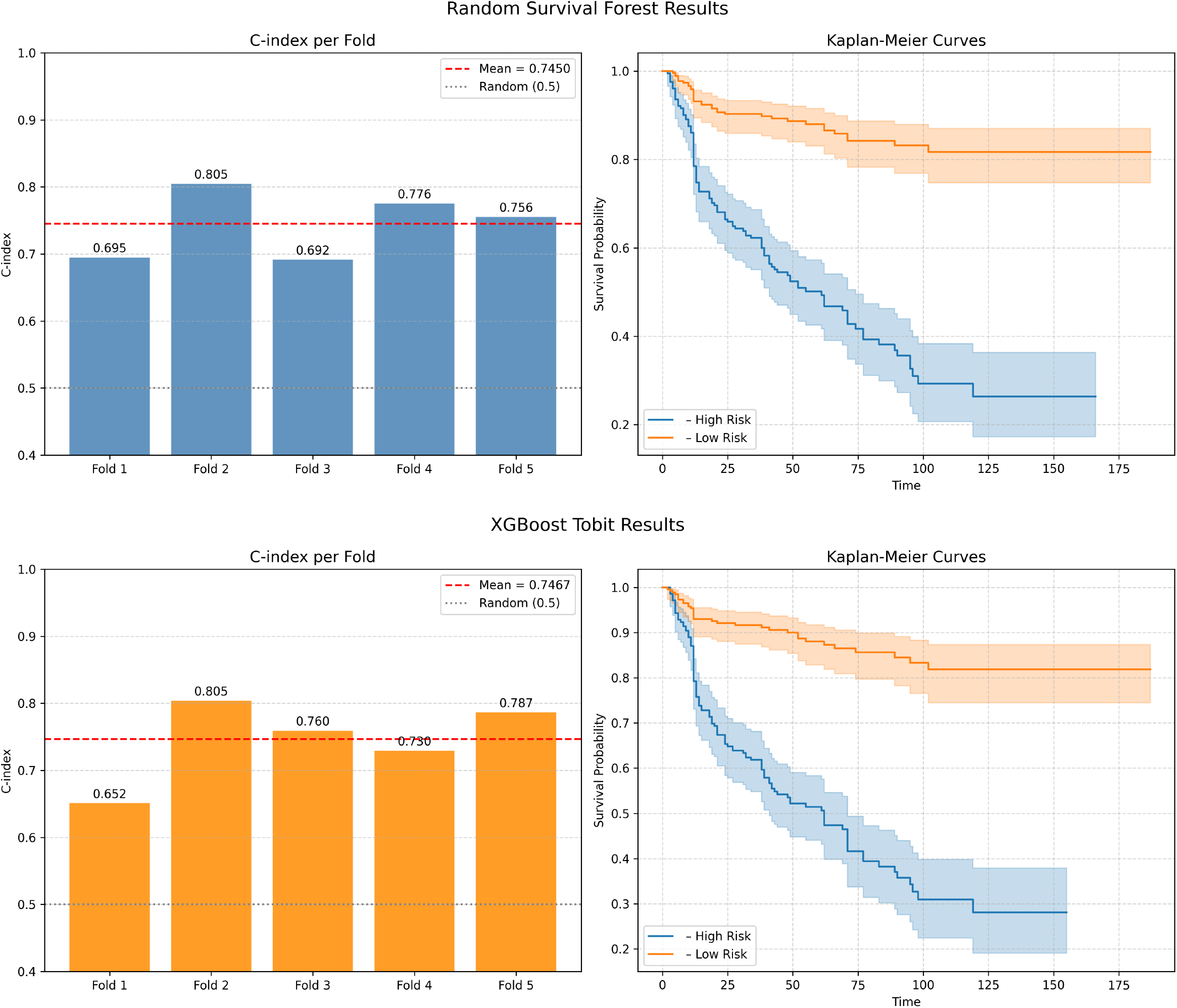
Survival analysis results for Random Survival Forest (top) and XGBoost Tobit (bottom). Left panels show the C-index per fold with 5-fold cross-validation; dashed red lines indicate the mean C-index. Right panels show Kaplan-Meier survival curves stratified by predicted risk group (high vs. low), with shaded regions indicating 95% confidence intervals.

**Figure S3.**
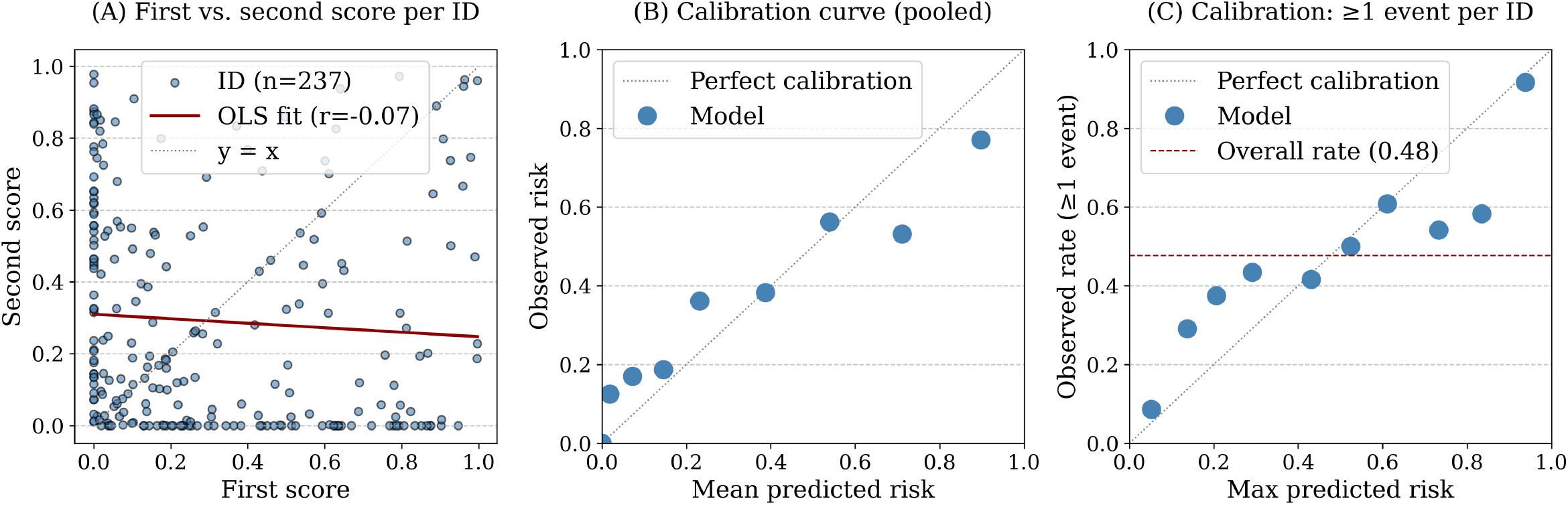
A) Score of first and second donor DQ heterodimers are slightly negatively correlated. B) Calibration plot - risk of XG- Boost score per allele, C) Calibration plot - risk of XGBoost score per donor.

## Literature cited

Aubert O, Loupy A, Hidalgo L, van Huyen JPD, Higgins S et al. 2017. Antibody-mediated rejection due to preexisting versus de novo donor-specific antibodies in kidney allograft recipients. Journal of the American Society of Nephrology. 28:1912–1923.

Barker DJ, Maccari G, Georgiou X, Cooper MA, Flicek P et al. 2023. The ipd-imgt/hla database. Nucleic acids research. 51:D1053–D1060.

Barnwal A, Cho H, Hocking T. 2022. Survival regression with accelerated failure time model in xgboost. Journal of Computational and Graphical Statistics. 31:1292–1302.

Callemeyn J, Lamarthée B, Koenig A, Koshy P, Thaunat O et al. 2022. Allorecognition and the spectrum of kidney transplant rejection. Kidney international. 101:692–710.

Chen T, Guestrin C. 2016. Xgboost: A scalable tree boosting system. In: . pp. 785–794.

Das R, Greenspan NS. 2025. Understanding hla-dq in renal transplantation: a mini-review. Frontiers in Immunology. 16:1525306.

Dayhoff M, Schwartz R, Orcutt B. 1978. 22 a model of evolutionary change in proteins. Atlas of protein sequence and structure. 5:345–352.

Demir Z, Raynaud M, Divard G, Louis K, Truchot A et al. 2025. Impact of hla evolutionary divergence and donor-recipient molecular mismatches on antibody-mediated rejection of kidney allografts. Nature Communications. 16:5692.

Duquesnoy RJ. 2008. Clinical usefulness of hlamatchmaker in hla epitope matching for organ transplantation. Current opinion in immunology. 20:594–601.

Duquesnoy RJ, Askar M. 2007. Hlamatchmaker: a molecularly based algorithm for histocompatibility determination. v. eplet matching for hla-dr, hla-dq, and hla-dp. Human immunology. 68:12–25.

Duquesnoy RJ, Marrari M. 2002. Hlamatchmaker: a molecularly based algorithm for histocompatibility determination. ii. verification of the algorithm and determination of the relative immunogenicity of amino acid triplet-defined epitopes. Human immunology. 63:353–363.

Duquesnoy RJ, Marrari M. 2009. Hlamatchmaker-based definition of structural human leukocyte antigen epitopes detected by alloantibodies. Current opinion in organ transplantation. 14:403–409.

Hafeez MS, Awais SB, Razvi M, Bangash MH, Hsiou DA et al. 2023. Hla mismatch is important for 20-year graft survival in kidney transplant patients. Transplant Immunology. 80:101861.

Hanley JA, McNeil BJ. 1982. The meaning and use of the area under a receiver operating characteristic (roc) curve. Radiology. 143:29–36.

Harrell FE, Califf RM, Pryor DB, Lee KL, Rosati RA. 1982. Evaluating the yield of medical tests. Jama. 247:2543–2546.

Ishwaran H, Kogalur UB, Blackstone EH, Lauer MS. 2008. Random survival forests. .

Ke G, Meng Q, Finley T, Wang T, Chen W et al. 2017. Lightgbm: A highly efficient gradient boosting decision tree. Advances in neural information processing systems. 30.

Kosmoliaptsis V, Gjorgjimajkoska O, Sharples LD, Chaudhry AN, Chatzizacharias N et al. 2014. Impact of donor mismatches at individual hla-a,-b,-c,-dr, and-dq loci on the development of hla-specific antibodies in patients listed for repeat renal transplantation. Kidney international. 86:1039–1048.

Kosmoliaptsis V, Mallon D, Chen Y, Bolton EM, Bradley JA et al. 2016. Alloantibody responses after renal transplant failure can be better predicted by donor–recipient hla amino acid sequence and physicochemical disparities than conventional hla matching. American journal of transplantation. 16:2139– 2147.

Kramer CS, Koster J, Haasnoot GW, Roelen DL, Claas FH et al. 2020. Hla-emma: a user-friendly tool to analyse hla class i and class ii compatibility on the amino acid level. Hla. 96:43–51.

Lundberg SM, Lee SI. 2017. A unified approach to interpreting model predictions. Advances in neural information processing systems. 30.

Maguire C, Crivello P, Fleischhauer K, Isaacson D, Casillas A et al. 2024. Qualitative, rather than quantitative, differences between hla-dq alleles affect hla-dq immunogenicity in organ transplantation. HLA. 103:e15455.

Mallon DH, Kling C, Robb M, Ellinghaus E, Bradley JA et al. 2018. Predicting humoral alloimmunity from differences in donor and recipient hla surface electrostatic potential. The Journal of Immunology. 201:3780–3792.

Pedregosa F, Varoquaux G, Gramfort A, Michel V, Thirion B et al. 2011. Scikit-learn: Machine learning in python. the Journal of machine Learning research. 12:2825–2830.

Saleem N, Das R, Tambur AR. 2022. Molecular histocompatibility beyond tears: The next generation version. Human Immunology. 83:233–240.

Senev A, Coemans M, Lerut E, Van Sandt V, Kerkhofs J et al. 2020. Eplet mismatch load and de novo occurrence of donor-specific anti-hla antibodies, rejection, and graft failure after kidney transplantation: an observational cohort study. Journal of the American Society of Nephrology. 31:2193–2204.

Shtossel O, Koren O, Louzoun Y. 2025. Left barrier loss for unbiased survival analysis prediction. IEEE Transactions on Pattern Analysis & Machine Intelligence. pp. 1–13.

Tambur AR. 2018. Hla-epitope matching or eplet risk stratification: the devil is in the details. Frontiers in Immunology. 9:2010.

Tambur AR, Campbell P, Chong AS, Feng S, Ford ML et al. 2020. Sensitization in transplantation: assessment of risk (star) 2019 working group meeting report. American Journal of Transplantation. 20:2652–2668.

Tambur AR, Campbell P, Claas FH, Feng S, Gebel HM et al. 2018. Sensitization in transplantation: assessment of risk (star) 2017 working group meeting report. American Journal of Transplantation. 18:1604–1614.

Tambur AR, Das R. 2023. Can we use eplets (or molecular) mismatch load analysis to improve organ allocation? the hope and the hype. Transplantation. 107:605–615.

Tamura K, Stecher G, Kumar S. 2021. Mega11: molecular evolutionary genetics analysis version 11. Molecular biology and evolution. 38:3022–3027.

Thaunat O, Koenig A, Leibler C, Grimbert P. 2016. Effect of immunosuppressive drugs on humoral allosensitization after kidney transplant. Journal of the American Society of Nephrology. 27:1890–1900.

Tian Y, Frischknecht L, Rössler F, Schachtner T, Nilsson J. 2025. De novo donor-specific hla antibody development after kidney transplantation is impacted by pirche ii score and recipient age. Frontiers in Immunology. 16:1508586.

Wiebe C, Kosmoliaptsis V, Pochinco D, Gibson IW, Ho J et al. 2019. Hla-dr/dq molecular mismatch: a prognostic biomarker for primary alloimmunity. American journal of transplantation. 19:1708–1719.

Wiebe C, Pochinco D, Blydt-Hansen T, Ho J, Birk P et al. 2013. Class ii hla epitope matching—a strategy to minimize de novo donor-specific antibody development and improve outcomes. American journal of transplantation. 13:3114–3122.

Wiebe C, Rush DN, Nevins TE, Birk PE, Blydt-Hansen T et al. 2017. Class ii eplet mismatch modulates tacrolimus trough levels required to prevent donor-specific antibody development. Journal of the American Society of Nephrology. 28:3353–3362.

